# Genetically Engineered DENV Produces Antigenically Distinct Mature Particles

**DOI:** 10.1101/2021.04.06.438747

**Authors:** Longping V. Tse, Rita M. Meganck, Stephanie Dong, Lily E. Adams, Laura J. White, Aravinda M. de Silva, Ralph S. Baric

## Abstract

Maturation of Dengue viruses (DENV) alters the structure, immunity and infectivity of the virion and highly mature particles represent the dominant form *in vivo*. The production of highly mature virions principally relies on the structure and function of the viral premature protein (prM) and its cleavage by the host protease furin. We developed a reliable clonal cell line which produces single-round mature DENVs without the need for DENV reverse genetics. More importantly, using protein engineering coupled with natural and directed evolution of the prM cleavage site, we engineered genetically stable mature DENVs without comprising viral yield and independent of cell, host, or passage. Using these complementary strategies to regulate maturation, we demonstrate that the resulting mature DENVs are antigenically distinct from their isogenic immature forms. Given the clinical importance of mature DENVs in immunity, our strategy provides a reliable strategy for the production of stable, high-titer mature candidate DENV live virus vaccines, genetically stabilized viruses for DENV maturation and immunity studies, and models for maturation-regulated experimental evolution in mammalian and invertebrate cells. Our data from directed-evolution across host species reveals distinct maturation-dependent selective pressures between mammalian and insect cells, which sheds light on the divergent evolutionary relationship of DENVs between its host and vector.

## Introduction

Mosquito-borne Dengue virus (DENV) is a major global public health threat causing ∼400 million new cases of dengue annually^1,2^. Although the majority of cases occur in tropical and subtropical areas where the mosquito vectors are most concentrated, global warming, travel, and globalization have contributed to the worldwide spread and intermixing of the four DENV serotypes^3^. Indeed, DENV infection has increased 30-fold between 1960 and 2010 with an upsurge of cases in the USA and Europe. A hallmark of DENV pathogenesis is the possibility for antibody dependent enhancement (ADE), which can progress to life threatening dengue hemorrhagic fever/dengue shock syndrome (DHF/DSS) upon secondary infection with a different serotype. So far, no antiviral treatments are available to treat DENV disease and the only approved vaccine, Dengvaxia, is not recommended for use in naïve populations^4,5^.

Proteolytic cleavage of viral membrane fusion proteins is a common strategy for temporal or spatial control of virus infection, ultimately affecting tropism and transmission^6,7^. The DENV virion structural proteins consists of capsid, E (envelope), and prM glycoproteins which undergo major conformational changes via the process of maturation. The most common depiction of DENV particles features the mature form, which is composed of 90 Envelope (E) homodimers lying flat in a “herringbone” structure and organized into a 50 nm icosahedral (T=pseudo 3) symmetry resembling other non-enveloped virions^8^. However, the virion is assembled in the ER as a non-infectious^9^, immature virion which adopts a completely distinct structure as a 60 nm “spikey” sphere with 60 three-fold spikes^10,11^. Each “spike” is composed of three E protein monomers elevated at a 27° upward angle with the fusion loop covered by prM proteins^11,12^. Maturation is a two-step process involving the proteolytic cleavage of prM by furin, a ubiquitously expressed serine protease with a preference for basic (positively charged) substrates, at the trans-Golgi network (TGN) followed by its release at neutral pH outside of the cell^11,13–15^. Cleavage of prM releases pr from the virion and triggers the E protein to rotate, ranging from ∼137° to ∼300°, to form the mature virion^10,16^.

While the maturation status of common laboratory DENV strains varies, one study showed that clinical isolates are typically more mature, arguing the clinical importance of mature DENVs^17^. Because the E protein undergoes major conformational changes during processing, mature and immature virions are predicted to present dramatically different combinations of antigenic structures and epitopes^18,19^. Further complicating the process, the conformational change is reversible (“breathing”) and patchy, as a single particle can adopt both mature and immature forms in different regions and at different times^20^. The biological functions and characteristics of these heterogeneous maturation forms remain largely unknown, but are thought to provide key evolutionary advantages in virus infection, immunity, and antigenic variation^21,22^.

Previous studies have shown that fully mature DENV can be generated in Vero cells overexpressing furin^23^. However, due to the polyclonal nature of the cells, viral yield as well as maturity depends on the cell passage and culture conditions. Furthermore, maturation phenotypes quickly switch from mature back to immature after a single replication, which limits assay usage to those not requiring viral replication. Importantly, maturation status can vary significantly between serotypes and genotypes, suggesting the presence of other, less understood, regulatory determinants^24^. In this study, we develop two complementary strategies, ectopic expression of furin in culture and virus genetic engineering, to produce mature virions across the four DENV serotypes. Additionally, we provide insight into the role of variation in the prM furin cleavage site as the major molecular determinant governing DENV maturation in vertebrate and invertebrate cells. Using protein engineering and directed-evolution, we generated high yields of mature DENV1, 2 and 4 using unmodified Vero cells. The current study advances our understanding of the biological and genetic processes of DENV maturation, develops novel tools and recombinant viruses, and provides further insight and essential tools for future investigations.

## Results

### DENV Maturity is Serotype Dependent

DENV maturation regulates virion infectivity and antigenicity and directly impacts antibody neutralization and potential vaccine efficacy. Since furin cleavage of the prM protein initiates the DENV maturation process, we hypothesized that furin cleavage efficiency is directly proportional to DENV maturation. Consequently, we compared the DENV1-4 prM cleavage site with other mosquito-borne Flaviviruses, highly pathogenic avian influenza virus (HPAI) and SARS-CoV-2. Sequence analyses suggested that all the DENV serotypes encoded a sub-optimal furin cleavage site (P4) R-X-K/R-R (P1) with negative modulators as indicated by an acidic residue at the P3 position (Fig. 1a). To analyze the functionality of the prM furin cleavage site in a more quantitative manner, we used the computational program PiTou, which combines machine learning and cumulative probability score function of known furin cleavages to calculate the logarithmic-odd probabilities of all the different viral furin cleavage sites^25^. These analyses provided strong predictive data that the DENV serotypes encode suboptimal furin sites (scores from 6.90 – 13.26) compared with other Flaviviruses (scores from 13.30 – 15.40) (Fig. 1a). We focused our studies on four prototypical wildtype (WT) DENV viruses including WestPac (DV1-WT), S16803 (DV2-WT), 3001 (DV3-WT) and Sri Lanka 92 (DV4-WT) isolates (Table 1). Using western blotting as a readout, we determined the relative maturity of each serotype by calculating the ratio of prM to E. Consistent with the hypothesis that prM cleavage is dependent on both local primary sequence and other distal and structural functions, PiTou predictions do not translate completely to the empirical maturation status of DENV. Relative maturity was clearly different between serotypes. In particular, serotypes encoding an Glutamic acid (E), but not Aspartic acid (D) at the P3 position (prM residue 89) are associated with more immature virion production in Vero cells, with DV2-WT virions containing the highest level of uncleaved prM, followed by DV4-WT, DV3-WT, and DV1-WT which has nearly undetectable levels of prM, and hence is more mature (Fig. 1b).

**Table 1:**
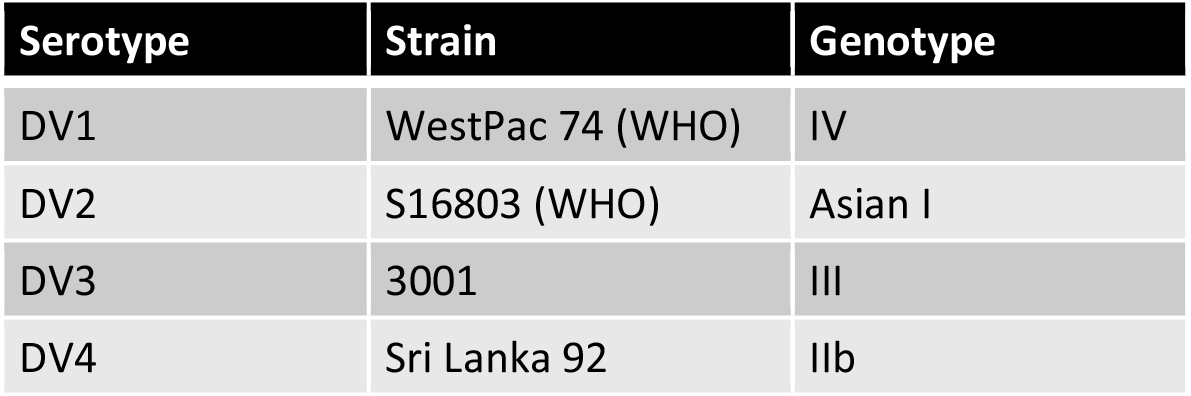
Prototypic WT DENV strains used in this study.

**Figure 1:**
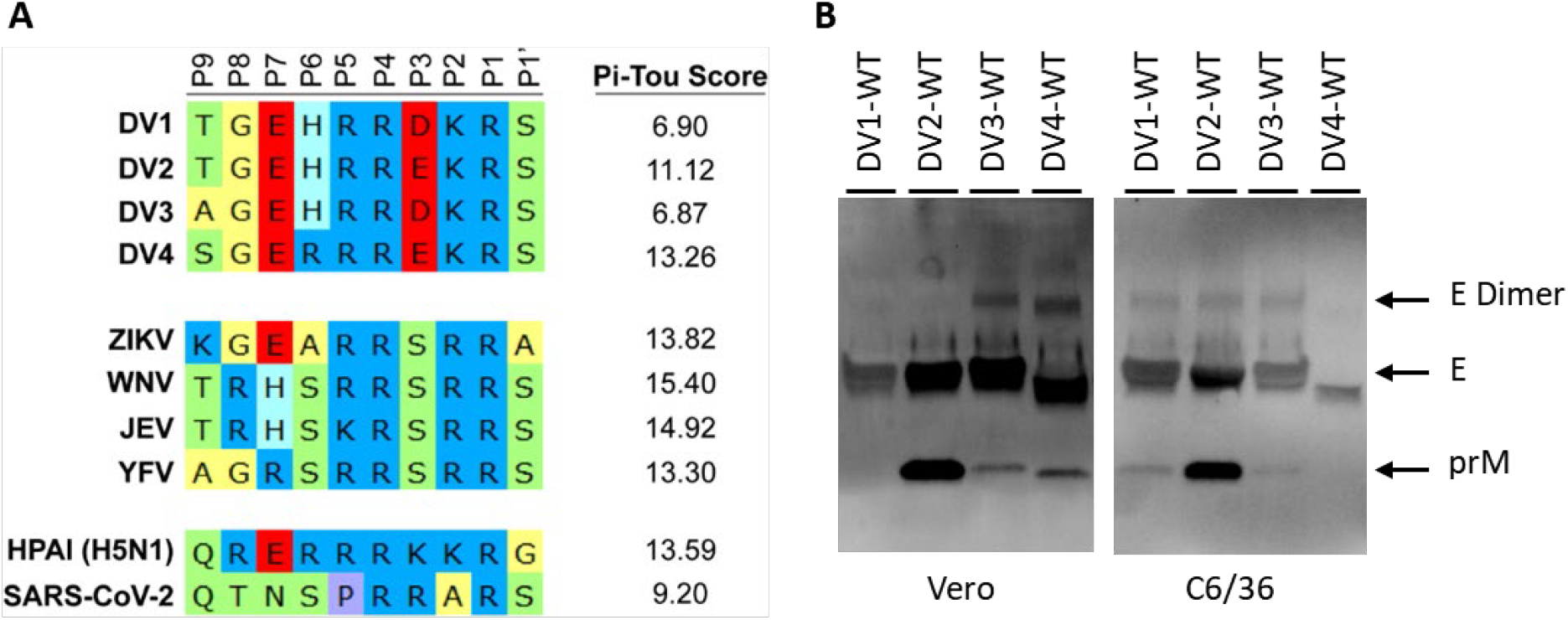
Furin cleavage site alignment and DENV maturation. (A) Amino acid sequence alignment of viral furin cleavage sites from position 9 (P9) to position 1 prime (P1’). Pi-Tou scores are the prediction of logarithmic-odd probabilities of all the different viral furin cleavage sites (higher value = better substrate for furin). DENV1, 2, 3 4, Zika virus (ZIKV), West Nile virus (WNV), Japanese encephalitis virus (JEV), yellow fever virus (YFV), highly pathogenic avian influenza virus (HPAI) and SARS-Coronavirus-2. (B) Representative western blot images of DENV 1-4 viral supernatants from Vero and C6/36 cells blotted with anti-Env and anti-prM antibodies.

### Optimized Clonal Vero-furin Cells Generate High Yield, Mature DENV

DENV maturation also depends on the producer cells; for instance, C6/36 grown DENVs show a different maturation profile, from DV2-WT (most immature) < DV1-WT = DV3-WT < DENV4 (most mature) (Fig. 1b). As reported previously^23^, fully mature DENV strains can be generated in Vero cells that overexpress furin. However, high level furin expression may negatively impact DENV virus production. Using the sleeping beauty transposon system^26^, we isolated two clonal lines with high (VF-Hi) or low (VF-Lo) levels of furin expression (Fig. 2a). Immunofluorescent staining and western blot analysis revealed different levels of furin expression in the trans-Golgi network (Fig. 2b and 2c). The growth kinetics of all four DENV serotypes were tested on both Vero-furin lines and compared to unmodified Vero cells (Fig. 2d – g). DV1-WT, DV2-WT and DV4-WT showed similar growth kinetics in all cell lines tested, while VF-Hi supported better DV3-WT growth (Fig. 2d – g). VF-Hi supports the production of fully mature DENV virions across all four serotypes (Fig. 2d – g). In agreement with the low furin expression level, VF-Lo phenocopied the DENV maturation status of unmodified Vero cells (Fig. 2d – g). Therefore, VF-Hi cells allow for high DENV yield in all serotypes, suggesting the furin expression level in VF-Hi is optimal for production of fully mature DENVs.

**Figure 2:**
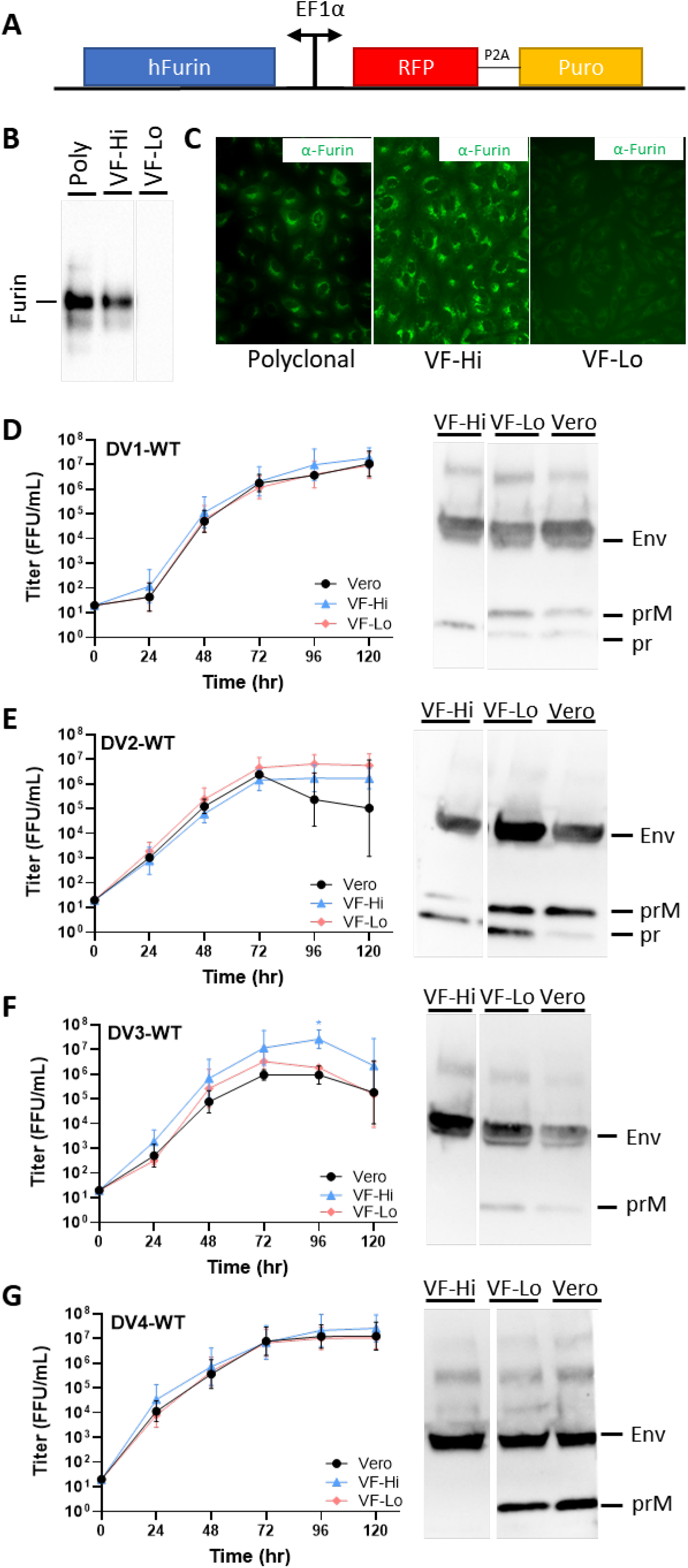
Growth kinetics and maturation status of Vero-furin grown DENVs. (A) Schematic of the Sleeping Beauty-based transposon cassette for ectopic expression of human furin (hFurin). A bi-directional EF1a promoter was used to drive the expression of hFurin and red-fluorescent protein (RFP) with a puromycin resistance gene (Puro) linked by a 2A self-cleaving peptide (P2A). (B) Western blot and (C) immunofluorescence images of polyclonal and clonal selected high (VF-Hi) and low (VF-Lo) expression hFurin Vero cells using anti-furin antibodies. Growth kinetics and degree of maturation of (D) DENV1-WT, (E) DENV2-WT, (F) DENV3-WT and (G) DENV4-WT in unmodified Vero cells (Black-Circle), VF-Hi cells (Blue-Triangle) and VF-Lo cells (Pink-Diamond). Cells were infected with DENV at MOI 0.01 – 0.05 for 120 hours. Supernatants were harvested at 120hpi and analyzed by western blot for DENV maturation using anti-Env and anti-prM antibodies. All assays were performed with at least two biological repeats with two technical replicates. Growth kinetics of DENV variants were compared to their corresponding wildtype using 2-way ANOVA multiple comparisons.

### Genetic Regulation of DENV1 and DENV4 Maturation Status

As an alternative to ectopic overexpression of furin which only generates mature virion for a single round of infection, we hypothesized that genetic modification of the prM furin cleavage site could also be used to optimize DENV maturation independence of cells or hosts. Using DV1-WT as a model, we introduced a mutation at the P3 position of the furin cleavage site and generated an isogenic strain, DV1-prM-D89K. The mutated cleavage site (HRR<u>K</u>KR|S) has a Pi-Tou score of 14.68 compared to the DV1-WT cleavage site (HRRDKR|S) with a Pi-Tou score of 6.90, predicting more optimal cleavage (Fig. 3a). DV1-WT and DV1-prM-D89K displayed no difference in virus growth kinetics in Vero (mammalian) and C6/36 (insect) cells (Fig. 3b). In both Vero and C6/36 cultures, DV1prM-D89K was more mature than DV1-WT, phenocopying the Vero-furin grown DV1-WT (Fig. 3c).

**Figure 3:**
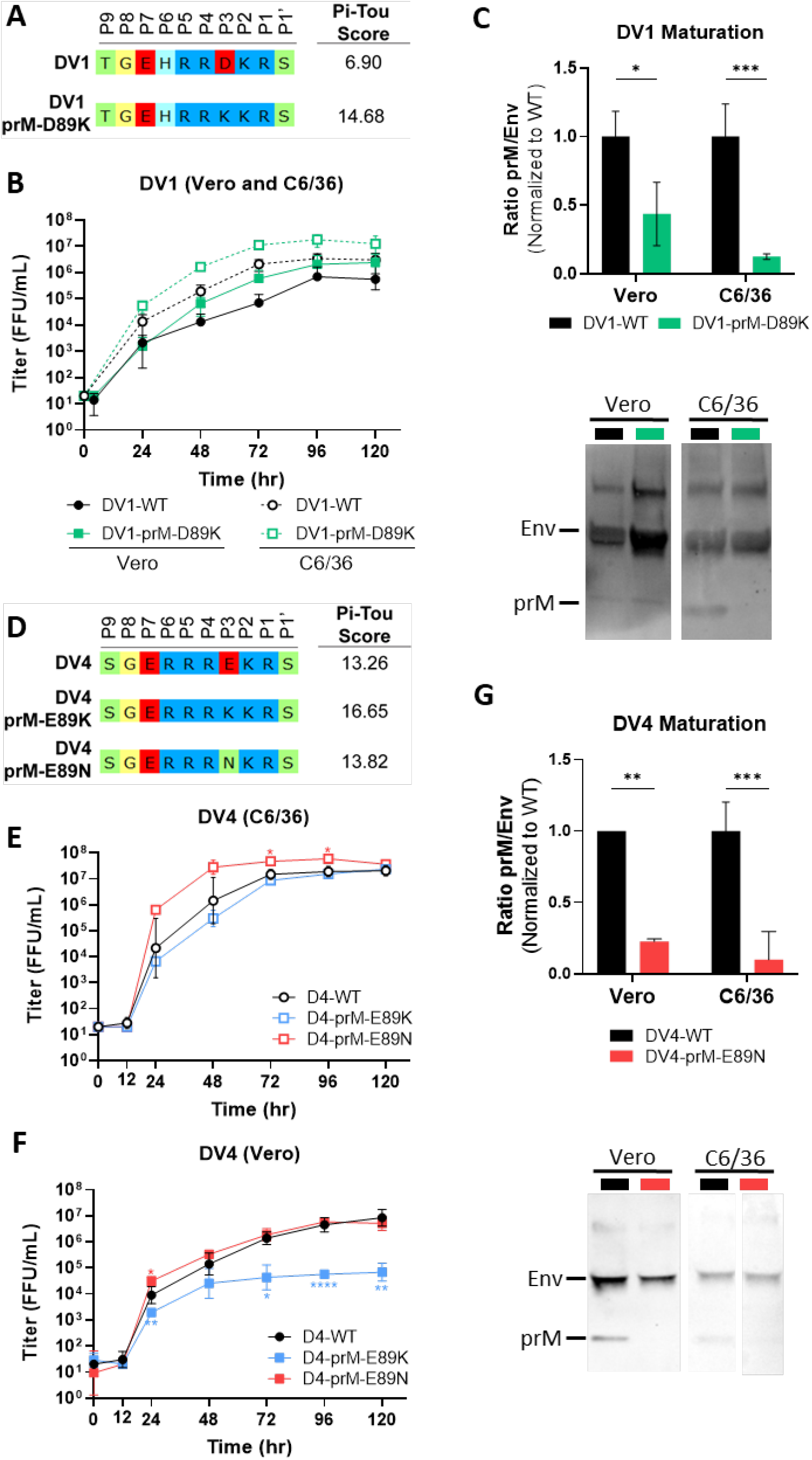
Generation of mature DENV1 and DENV4 via genetic modification. (A) Sequence alignment and Pi-Tou scores of DV1-WT and DV1-prM-D89K. (B) Growth kinetics of DV1-WT and DV1-prM-D89K in Vero and C6/36 cells. (C) Representative western blot image (bottom) of DV1-WT and DV1-prM-D89K viral supernatants blotted with anti-Env and anti-prM antibodies, and quantification (top) of viral maturation (prM/Env) normalized to DV1-WT (lower value = more mature). (D) Sequence alignment and Pi-Tou scores of DV4-WT, DV4-prM-E89K and DV4-prM-E89N. Growth kinetics of DV4-WT, DV4-prM-E89K and DV4-prM-E89N in (E) C6/36 and (F) Vero cells. (G) Representative western blot image (bottom) of DV4-WT, DV4-prM-E89K and DV4-prM-E89N viral supernatants blotted with anti-Env and anti-prM antibodies and quantification (top) of viral maturation (prM/Env) normalized to DV4-WT (lower value = more mature). Growth kinetics and maturation of DENV variants were compared to their corresponding wildtype using 2-way ANOVA multiple comparisons.

To understand if the furin cleavage site mutation is portable across serotypes, we introduced a similar mutation on the DV4-WT backbone, generating the isogenic strain DV4-prM-E89K (Fig. 3d). While we successfully generated a pure population of DV4-prM-E89K in C6/36 cells, a spontaneous mutation, K89N, rapidly emerged and gave rise to a new evolved DV4-prM-E89N variant in Vero cells by passage 2 (Fig. S1a). By the 5^th^ passage, the DV4-prM-E89N variant represented 100% of the viral population (Fig. S1a), supporting the notion that viruses encoding the E89K mutation were less fit than those encoding the E89K mutation in Vero cells. Growth kinetics of DV4-prM-E89K and DV4-prM-E89N on C6/36 cells are comparable to DV4-WT (Fig. 3e). However, the DV4-prM-E89K variant displayed a robust 2-log growth defect compared to DV4-prM-E89N and DV4-WT on Vero cells (Fig. 3f). When grown at 32°C, the growth defect of DV4-prM-E89K was alleviated (Fig. S1c). The maturation status of the two variants were tested and compared to DV4-WT. DV4-prM-E89N is more mature than DV4-WT in both Vero and C6/36 cells (Fig. 3g). No prM can be detected in DV4-prM-E89K; due to the low virus yield in Vero cells, the data suggest that either DV4-prM-E89K is fully mature or the protein input is below detection limit (Fig. S1b). As calculated by Pi-Tou, DV4 has the highest furin cleavage score among the DENV serotypes at 13.26. The point mutation prM-E89K increases the score to 16.65 (the highest score observed), while DV4-prM-E89N has a Pi-Tou score of 13.82 (Fig. 3d). In DENV4, it seems that a “super-optimal” furin cleavage site may negatively impact DENV growth in Vero cells. The data suggest a delicate balance likely exists between virion maturation, furin cleavage site efficiency, and viral fitness in different serotypes.

### Directed Evolution Reveals High Levels of Plasticity in DENV2 prM Cleavage Site

Based on the spontaneous K89N mutation in DV4, we hypothesized the prM cleavage site has high plasticity, suggesting the existence of a “Goldilocks Zone” for efficient *in vitro* growth. We utilized saturation mutagenesis and directed-evolution to simultaneously screen thousands of DENV2 prM cleavage site variants for efficient growth in tissue culture. We generated a DENV2 viral library in which four positions, P3, P5, P6, and P7, of the prM cleavage site were randomly mutated, preserving the core furin cleavage site (Fig. 4a). The library was propagated three times in either Vero or C6/36 cells, and each passage of the virus were deep sequenced along with the plasmid library (Fig. 4a). The theoretical amino acid diversity of the library is 160,000 variants (ignoring stop codons), which was represented in the plasmid library (Table 2). As expected, viral diversity rapidly drops after one passage, to 0.7% (1148 unique variants) and 16.2% (25942 unique variants) of the theoretical maximum in Vero and C6/C6 respectively, further diminished after each passage (Table 2). The large number of viable DENV2 variants in both cells indicates a high degree of plasticity within the prM cleavage site in culture (Table 2). Importantly, C6/36 cells were more tolerant to prM cleavage site variations than Vero, suggesting a higher selective pressure exerted by mammalian cells. After three rounds of selection in C6/36 and Vero cells, two different dominant variants, TG<u>RAQ</u>R<u>Y</u>KR|S (DV2-C1) and TG<u>AGR</u>R<u>S</u>KR|S (DV2-V1), emerged, representing almost 50% of their respective viral populations (Fig 4b and 4c). While the DV2-WT cleavage site has a Pi-Tou score of 11.12, the Vero-selected cleavage site score increased to 14.39. Surprisingly, the DV2-C1 cleavage site scored at 7.76, a much lower score than DV2-WT (Fig. 4d). We plotted the PiTou score distribution of the top 50 ranked variants in C6/36 and Vero cells, with peaks at 7.7 and 14.9, respectively (Fig. 4e). We also plotted the Pi-Tou scores of the top 50 sequences from passage 1 that were extinct by passage 3. Although there was no distinct peak of deselection in C6/36 cells, a distinct peak of Pi-Tou scores at 13.9 were observed in the Vero-selected extinct population (Fig. 4e). The sequences, counts, and Pi-Tou scores of the top 50 enriched and deselected cleavage sites are summarized in Table S1 and S2. Due to founder effects in directed-evolution experiments, there is only one sequence shared between the top 50 variants evolved from Vero and C6/36 cells after three passages. Additionally, some variants with high Pi-Tou scores are rapidly deselected in both Vero and C6/36 cells, suggesting that the furin cleavage site sequence plays multiple roles in viral fitness (Table S2). The difference in scores between the two cell lines and the leveling effect of the lower ranked variants highlighted the differential fitness requirements of DENV2 between insect and mammalian cells.

**Table 2:** Summary of plasmid and viral passages diversities of DV2 directed-evolution.

|  | C6/36 Evolved |  | Vero Evolved |  |
| --- | --- | --- | --- | --- |
|  | Unique Sequences | % Maximum | Unique Sequences | % Maximum |
| Plasmid | 164569 | 102.86* | 164569 | 102.86* |
| P1 | 25942 | 16.21 | 1148 | 0.72 |
| P2 | 14119 | 8.82 | 719 | 0.45 |
| P3 | 6026 | 3.77 | 683 | 0.43 |

\* Plasmid library contains some sequences with stop codons.

**Figure 4:**
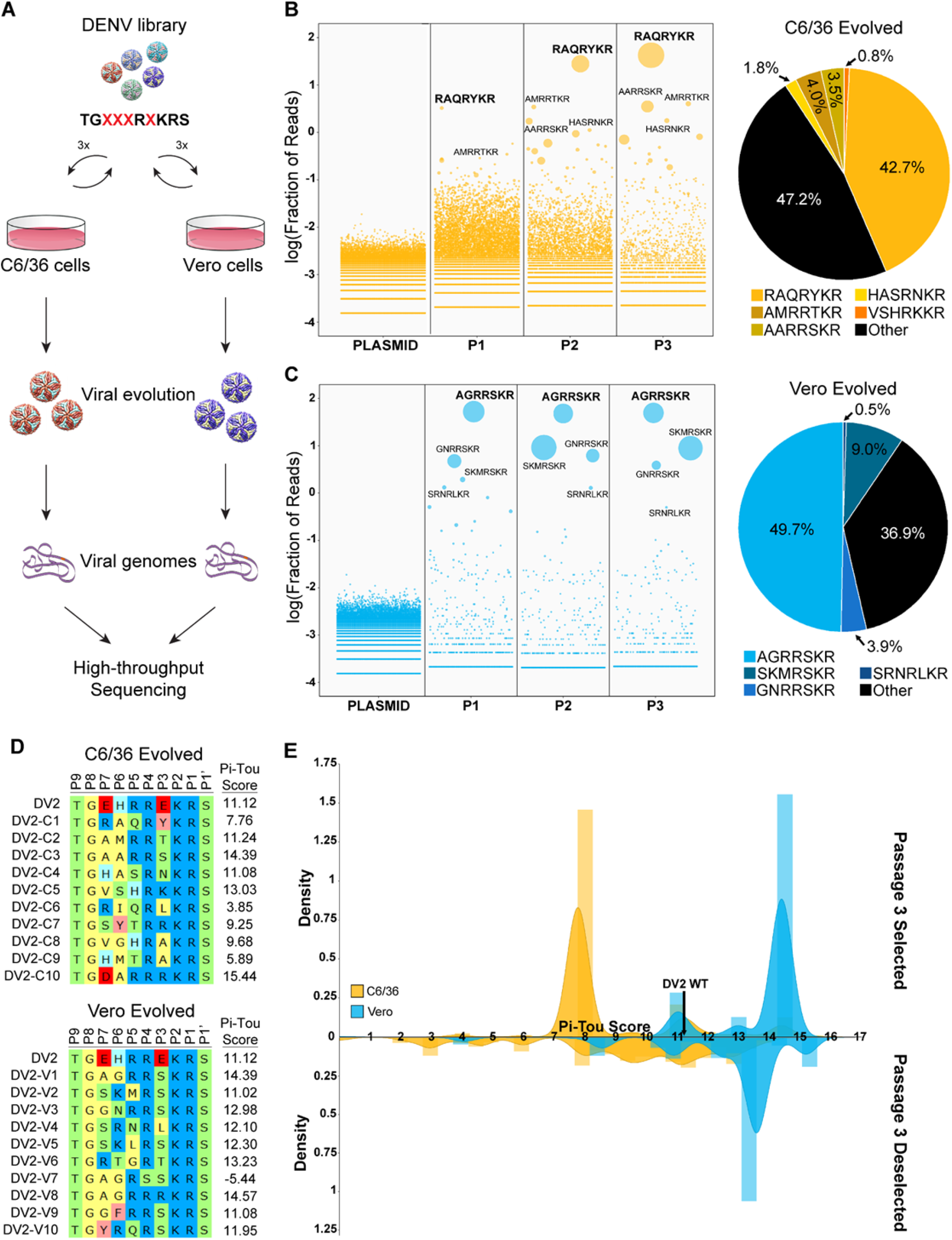
Directed-evolution of DENV2 prM cleavage site in Vero and C6/36 cells. (A) Schematic of directed-evolution from library generation to high-throughput sequencing. Enrichment plot of prM cleavage site sequences from plasmid library to viral population at the 3^rd^ passage (P3) and the proportion as well as sequence of the Top 5 enriched sequences in (B) C6/36 (yellow) and (C) Vero cells (Cyan). (D) Sequences and Pi-Tou scores of the Top 10 enriched prM cleavage sites from C6/36 and Vero cells. (E) Top: Distribution plot of Pi-Tou scores from the top 50 enriched prM cleavage sites in the 3^rd^ passage of C6/36 (yellow) and Vero cells (Cyan). PiTou score of DV2-WT is marked at 11.12 with a dark line. Bottom: Distribution plot of Pi-Tou scores of the top 50 variants present at the 1^st^ passage but lost in the 3^rd^ passage of C6/36 (yellow) and Vero cells (Cyan).

The top ranked evolved variants, DV2-V1 and DV2-C1, were re-derived via reverse genetics for further characterization. We also included a DV2-prM-E89K variant similar to the original DV1 mutation as comparison (Fig. 5c). While the DV2-prM-E89K variant has slightly reduced growth in Vero cells compared to DV2-WT, both DV2-V1 and DV2-C1 grow better than DV2-WT in Vero, with a drop in titer in C6/36 cells at 96 to 120 hpi (Fig. 5a and 5b). In Vero cells, DV2-prM-E89K and DV2-V1 are almost fully mature while DV2-C1 is only 30% more mature than DV2 WT (Fig. 5d). When the viruses are grown in C6/36, all the variants are 60 – 70% more mature than DV2 WT (Fig. 5d).

**Figure 5:**
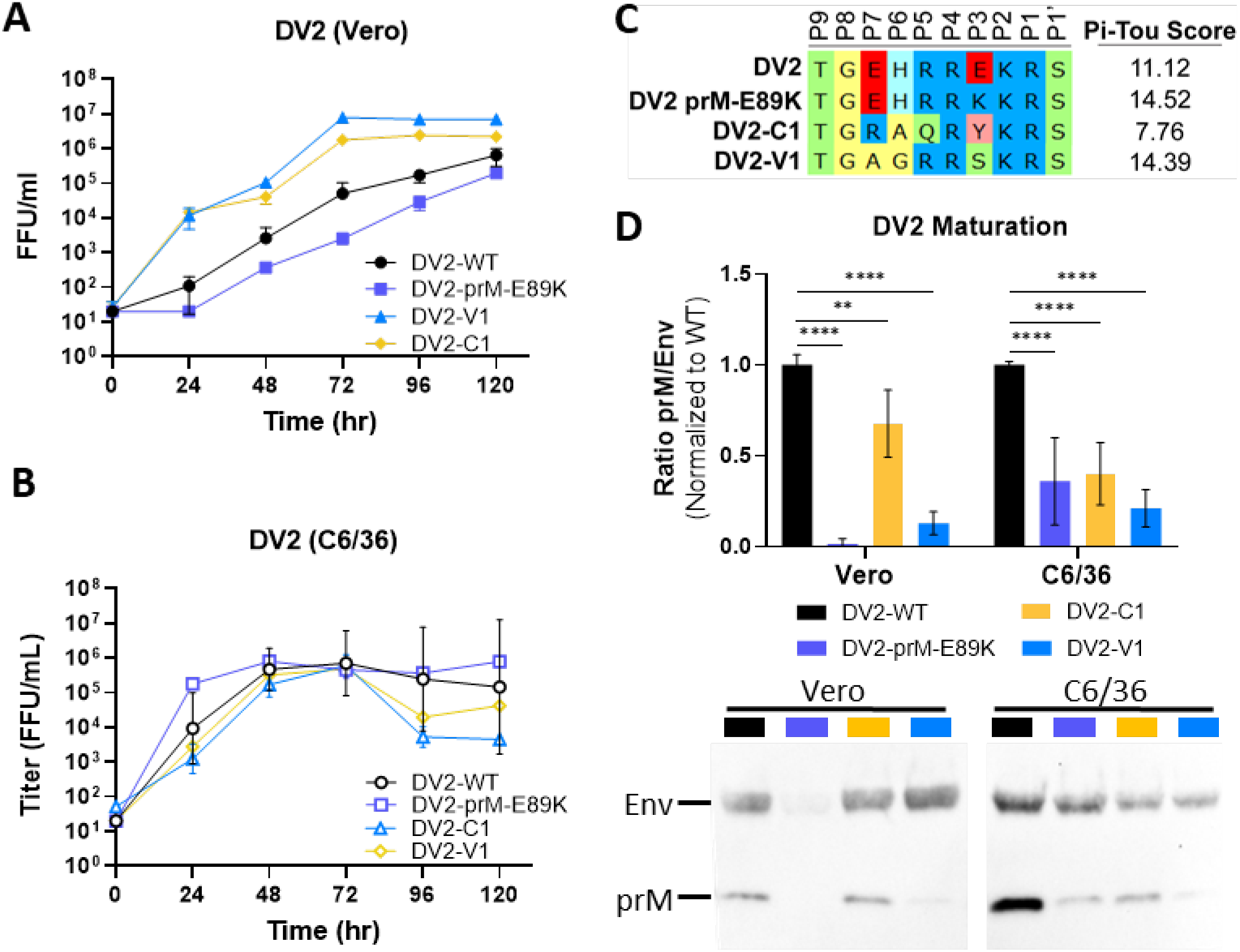
Generation of mature DENV2 via directed-evolution. Growth kinetics of DV2-WT, DV2-prM-E89K, DV2-C1 and DV2-V1 in (A) Vero and (B) C6/36 cells. (C) Sequence alignment and PiTou scores of DV2-WT, DV2-prM-E89K, DV2-C1 and DV2-V1 (D) Representative western blot image (bottom) of DV2-WT, DV2-prM-E89K, DV2-C1, and DV2-V1 viral supernatants blotted with anti-Env and anti-prM antibodies, and quantification (top) of viral maturation (prM/Env) normalized to DV2-WT (lower value = more mature). Growth kinetics and maturation of DV2 variants were compared to DV2-WT using 2-way ANOVA multiple comparisons.

### Impact of Maturation Status on DENV Epitope Presentation and Antigenicity

Given the ability to generate fully mature DENVs, we next evaluated the impact of maturation status on antigenicity. We selected several monoclonal antibodies targeting different regions of the DENV E glycoprotein, including C10 (Envelope-Dimer-Epitope 1)^27^, B7 (Envelope-Dimer-Epitope 2)^27^, 1C19 (BC loop)^28^ and 1M7 (fusion loop)^28^. As expected, Ab epitopes that are not maturation dependent are preserved, as evidenced by antibodies such as C10, B7, and 1C19 which showed no difference in <u>F</u>oci <u>R</u>eduction <u>N</u>eutralization <u>T</u>iter <u>50</u> values (FRNT50) (Fig. 6a – c). However, the fusion loop targeting antibody 1M7 showed significantly different FRNT50 values between fully mature and less mature DENVs in DENV1 and 4, but not in DENV2 (Fig. 6a – c). For DENV4, we also tested polyclonal sera from patients 180 days post DENV4 vaccination or naturally infected patients from a traveler cohort. Polyclonal serum contains a mixture of antibodies which may or may not be affected by virion maturation status. Unsurprisingly, FRNT_50_ of polyclonal serum was equivalent for fully mature and partially mature DENV4 (Fig. 6d).

**Figure 6:**
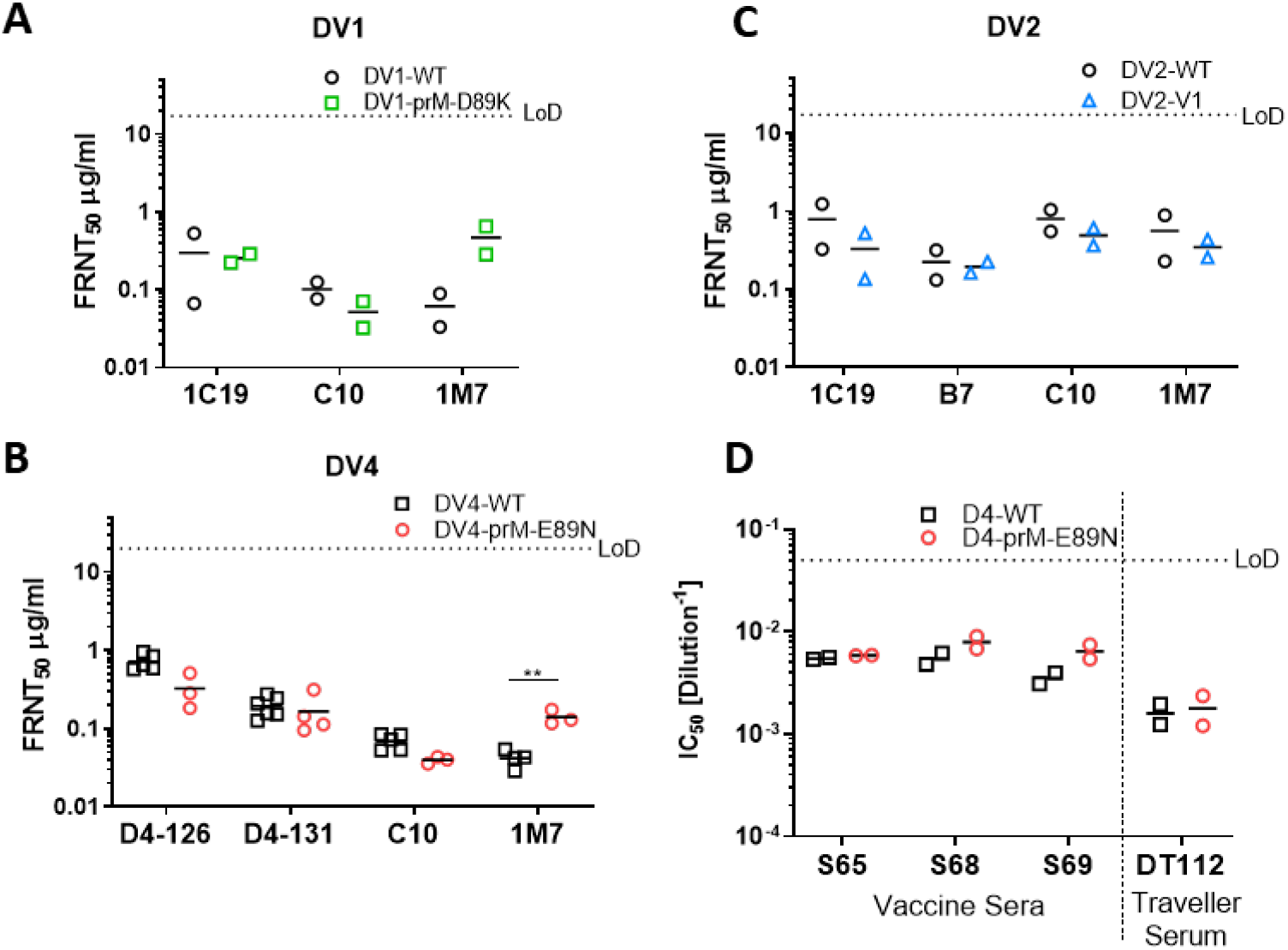
Antigenic profile of mature DENV. (A) FRNT_50_ of DV1-WT and DV1-prM-D89K against 1C19, EDE1-C10 (C10) and 1M7. (B) FRNT_50_ of DV4-WT and DV4-prM-E89N against D4-126, D4-131, C10 and 1M7. (C) FRNT_50_ of DV2-WT and DV2-V1 against 1C19, EDE2-B7 (B7), C10 and 1M7. (D) FRNT_50_ of vaccine sera and traveler serum against DV4 and DV4-prM-E89N.

## Discussion

In this report, we provided two methods to produce fully mature DENVs, and demonstrated that the prM furin cleavage site is the main determinant of maturation. The minimal furin cleavage site only requires (P4) R-X-K/R-R (P1), but cleavage efficiency greatly depends on positions P7, P6, P5, P3 and P1’ – P4’^29–31^. Algorithms such as Pi-Tou (used here) and ProP utilize machine learning to predict furin cleavage site efficiency while Pi-Tou also account for cumulative probability score function of known furin cleavage^25,32^. It must be noted that both programs focus on mammalian furin rather than invertebrate furin proteases; human furin and *Aedes aegypti* furin-like-proteases only share ∼40% sequence identity (Fig. S2) and Drosophila furin has been shown to have different substrate preferences^33^. Perhaps unsurprisingly, Pi-Tou predictions do not correlate fully with DENV maturation, as other determinants such as cleavage site accessibility, protein structure, stability, and the stem region of prM can also affect maturation in DENV and other viruses^34–36^. In DENV4, the degree of maturation differs among genotypes with identical prM proteins, suggesting contributions by an envelope-dependent maturation determinant as well^24^. A previous study generated mature dengue virus-like-particles (VLPs) by modifying the prM cleavage site for optimal cleavage^37^. In the current work, modification of the furin cleavage site and removal of the acidic P3 residues generated fully mature live DENV1, 2, and 4. However, unlike with Dengue-VLPs, experiments with authentic virus exposed the fitness cost of these mutations. The large growth defect of the DV4-prM-E89K variant was alleviated by reducing the temperature from 37°C to 32°C during virus production, indicating that the mutation impacts stability^18^, while the spontaneous mutation K89N restored viral fitness. These results hint at the presence of a maturity-stability balance in DENV, and the unfavorable acidic residues at P3 may play a regulatory role.

Using directed-evolution, we tested the fitness of thousands of DENV2 prM cleavage site variants, revealing high sequence plasticity. Predicted cleavage efficiency varied greatly between Vero- and C6/36-selected variants, further indicating differences in substrate preference of mammalian and insect furin which warrant further investigation. We observed many more viable variants in C6/36 cells compared to Vero cells, which may indicate a higher tolerance of mutation in furin sites in insect cells which could drive viral diversity and emergence in nature. However, we cannot rule out that the greater number of viable variants is an artifact of greater efficiency of the DENV reverse genetics system in C6/36 cells. Both of the top Vero- and C6/36-selected variants displayed enhanced growth kinetics and a slight increase in peak titer in Vero cells, indicating tissue culture adaption of the prM cleavage site; this advantage may not be reflected in natural infections or *in vivo*. Alignment of DENV prM sequences indicates that the furin cleavage site is extremely conserved in nature, despite high experimental plasticity. This discrepancy suggests either an unknown advantage of the WT cleavage site or a bottleneck effect in nature. Nevertheless, both variants from our directed evolution experiment are more mature than DV2-WT, suggesting selection for mature DENV2 *in vitro* in both mammalian and inset cells. Future experiments could focus on *in vivo* evolution of the genetic pool, using Aedes aegypti mosquitoes.

Based on a small cohort of monoclonal Abs and anti-DENV serum, our genetically modified mature DENVs show similar neutralization profiles to wildtype against antibodies targeting maturation independent epitopes (such as EDE1, EDE2 and BC loop epitopes), suggesting mutations at the prM cleavage site do not affect the overall viral protein structure. However, maturation dependent epitopes present only in one form of the virus, such as the fusion loop^17^, show a different neutralization profile against our mature DENVs. DENV2 has been shown to be very flexible, and “breathing” could account for the insensitivity of 1M7 neutralization to maturation status^38,39^. Our results support earlier studies showing differences in antigenicity between mature and immature DENVs using furin overexpression cells^18^, while providing new opportunities for studying the role of maturation in antigenicity, vaccine design, and *in vivo* replication and pathogenesis. Recent studies have demonstrated a potential disconnect between neutralizing antibody correlates of protective immunity in vaccine recipients^40^. Our data are consistent with earlier studies showing that the maturation status of DENV particles could have major implications for neutralization assay outcomes and result in bias during the determination of “correlates of protection” for vaccine studies^17,20,41^. These findings reinforce the importance of monitoring DENV maturation status in vaccine development, and our engineered strains provide a universal way to control DENV maturation for live-attenuated vaccine candidates independent of cell and host.

Like many studies, our report generated additional questions. Biologically, does DENV maturation play a more critical role than simply preventing premature fusion during production? Could maturation also play a role in vector-to-host or host-to-vector transmission? Is fully mature DENV advantageous or deleterious in mosquitoes and mammals? What determinants outside of the primary cleavage site sequence regulate maturation efficiency? Will biologically stabilized virions drive selection of unique subsets of neutralizing antibodies after infection? Clinically, the antigenic differences between mature and immature DENV require more comprehensive investigation. Furthermore, a new class of vaccines could be imagined based on stabilized mature particles which elicit maturation discriminatory antibodies. Given the clinical relevance and enigmatic nature of DENV maturation, our study adds to understanding of DENV maturation control and provides essential tools for future investigations.

## Materials and Methods

### Cells, plasmids and viruses

Mosquito (*Aedes albopictus*) C6/36 cells (ATCC# CRL-1660) were maintained in minimum essential medium (MEM) (Gibco) media supplemented with 5% fetal bovine serum (FBS) (HyClone), 100 U/mL penicillin and 100 mg/mL penicillin/streptomycin (P/S) (Gibco), 0.1 mM nonessential amino acids (NEAA) (Gibco), HEPES (Gibco) and 2 mM glutaMAX (Gibco) and incubated in the presence of 5% CO_2_ at 32°C. Vero (ATCC# CCL-81), VF-Hi and VF-Lo (generated from this study) were maintained in Dulbecco′s Modified Eagle′s Medium (DMEM) (Gibco) supplemented with 10% FBS, P/S, NEAA and HEPES and incubate in 5% CO_2_ at 37°C. DENV variants were generated by site-directed mutagenesis using Q5 High-fidelity DNA polymerase (NEB) followed by DENV reverse genetics (see below). The Env and prM of all DENV variants were sequence confirmed. DV1, 2, 3 and 4-WT viruses are grow in C6/36 or Vero cells maintained in infection media. C6/36 infection media contains Opti-MEM (Gibco) supplemented with 2% FBS, 1% P/S, 0.1 mM NEAA, 1% HEPES and 2 mM glutaMAX. Vero infection media is the same as the growth media except with 2% FBS supplement.

### DENV reverse genetics

Recombinant viruses were constructed using a four-plasmid cloning strategy as described previously^42^. The DENV genome was divided into four fragments (A−D fragment) and subcloned into four separate plasmids. A T7 promoter was introduced into the 5′ end of the A fragment, and unique type IIS restriction endonuclease cleavage sites are introduced into the 5′ and 3′ ends of each fragment to allow for systematic assembly into a full-length cDNA from which the full-length RNA transcripts can be derived. Plasmid DNA was grown in Top10 chemical component cells (ThermoFisher), digested with the corresponding enzymes, gel purified, and ligated together with T4 DNA ligase (NEB). Ligation products were purified by chloroform extraction. The purified ligation product was used as a template for *in-vitro* transcription to generate infectious genome-length capped viral RNA transcripts using T7 RNA polymerase (ThermoFisher). RNA was electroporated into either C6/36 or Vero cells. Cell culture supernatant containing virus was harvested 4 – 5 days post-electroporation as passage zero. During the subsequent passages following infection, the cells were grown in infection media.

### Stable cell line generation, VF-Hi and VF-Lo

Human furin was cloned in the sleeping beauty transposon plasmid^26^ pSB-bi-RP (Addgene #60513), transfected along with transposase, pCMV(CAT)T7-SB100 (Addgene #34879) into Vero cell using PEI Max (MW 40,000) (Polysciences) and selected with 2.5 µg/ml Puromycin (Gibco). Clonal cell lines were generated through limited dilution of the polyclonal cell line on a 96-well plate at the concentration of 0.3 cell/well.

### DENV growth kinetic and quantification

500,000 Vero or C6/36 cells were seeded in each well of a 6-well plate 1 day prior infection. Cells were infected with DENV at 0.05 to 0.1 M.O.I. assuming 1×10^6^ cells on the day of infection. Cells were washed 3 times with PBS and replenished with 3 mL of infection media after 1 hour of inoculation at 37°C in 5% CO_2_ incubator. 300 µl of viral supernatant was collected and fresh media was replenished at 0, 24, 48, 72, 96 and 120 hpi and stored at -80°C. Titer of the viral supernatant was determined using a standard DENV foci forming assay. In brief, Vero cells were seeded at 2×10^4^ cells/well in a 96-well plate. 50 µl of serial diluted viral supernatant were added to each well and incubated for 1h at 37°C in 5% CO_2_ incubator. 125 µl of overlay (Opti-MEM + 5% methyl cellulose + NEAA + P/S) was added to each well and incubated for 48h at 37°C + 5% CO_2_. Each well was rinsed 3 times with PBS and fixed with 10% formalin in PBS for staining. Vero cells were blocked in permeabilization buffer (eBioscience) with 5% non-fat dried milk. Two primary antibodies, anti-prM mAb 2H2 and anti-Env mAb 4G2, from non-purified hybridoma supernatant were used at 1:500 dilution in blocking buffer. Goat anti-mouse secondary conjugated with horseradish peroxidase (HRP) (SeraCare’s KPL) were diluted at 1:1000 in blocking buffer. Foci were developed using TrueBlue HRP substrate (SeraCare’s KPL) and counted using an automated Immunospot Analyzer instrument (Cellular Technology Limited). All experiments were performed independently a minimum of 3 times.

### Immunostaining and western blotting for human furin

Cells were fixed in 10% formalin in PBS and permeabilized with permeabilization buffer (eBioscience). Rabbit anti-furin (Thermo, PA1-062, 1:1000) was used as primary antibody. Goat anti-rabbit Alexa488 (Invitrogen, 1:2000) as secondary antibody. For western blotting, cell were lysed in 1% TritonX100, 100 mM Tris, 2M NaCl and 100 mM EDTA. Cell lysates were run in SDS-PAGE and blotted onto PVDF membrane. Furin bands were detected using rabbit anti-furin polyclonal at 1:1000 and Goat anti-rabbit HRP (Invitrogen, 1:5000) was used as secondary antibody.

### Western Blotting for DENV maturation

Viral stocks or supernatant from DENV growth curves at 120hpi were diluted with 4x Laemmli Sample Buffer (Bio-Rad) and boiled at 95°C for 5 minutes. Following SDS-PAGE electrophoresis, proteins were transferred to PVDF membrane and blocked in blocking buffer consist of 3% non-fat milk in PBS + 0.05% Tween-20 (PBS-T). The membrane was incubated with polyclonal rabbit anti-prM (1:1000, Invitrogen, Cat. #PA5-34966) and purified human anti-Env (fusion loop) 1M7 (2µg/ml) in 2% BSA + PBS-T solution for 1h at 37°C. The primary antigen-antibody complex was detected by incubating the blot with goat anti-rabbit IgG HRP (1:10000, Jackson-ImmunoLab) and sheep anti-human IgG HRP (1:5000, GE Healthcare) in 3% milk in PBS-T, for 1h at room temperature. Membranes were developed by Supersignal West Pico PLUS Chemiluminescent Substrate (ThermoFisher). Western blot images were captured with iBright FL1500 imaging system (Invitrogen). The pixel intensity of individual bands was measured using ImageJ, and relative maturation was calculated by using the following equation: (prM_Exp_/Env_Exp_)/(prM_WT_/Env_WT_). All experiments were performed independently a minimum of 3 times.

### Foci reduction neutralization titer assay (FRNT Assay)

FRNT assays were performed on Vero cells as has been described previously^43^. Briefly, 2×10^4^ Vero cells were seeded in a 96-well plate. Antiserum or mAbs were serially diluted and mixed with DENV viruses (80 – 100 FFU/well) at a 1:1 volume ratio and incubated at 37°C for 1h without the cells. The mixture was transferred to the 96-well plate with Vero cells and incubated at 37°C for 1h. The plate is subsequently overlaid with overlay medium (see above). Viral foci were stained and counted as described above. Data were fitted with variable slope sigmoidal dose-response curves and FRNT_50_ were calculated with top or bottom restraints of 100 and 0, respectively. All experiments were performed independently at least 2 times, due to limited amounts of human serum.

### DENV2 library generation and directed-evolution

DENV prM libraries were engineered through saturation mutagenesis on amino acid residues P3, 5, 6 and 7 of the DENV furin cleavage site based on previously published protocol^44^. In brief, degenerate NNK oligos (Integrated DNA Technologies) were used to amplify the prM region to generate a library with mutated prM DNA fragments. To limit bias and ensure accuracy, Q5 high fidelity polymerase (NEB) was used and limited to <18 cycles of amplification. The DNA library was cloned into the DENV reverse genetics system plasmid A to create a plasmid library by standard restriction digestion. Ligation reactions were then concentrated and purified by ethanol precipitation. Purified ligation products were electroporated into DH10B ElectroMax cells (Invitrogen) and directly plated on multiple 5,245-mm^2^ bioassay dishes (Corning) to avoid bias from bacterial suspension cultures. Colonies were pooled and purified using a Maxiprep Kit (Qiagen). The plasmid library was used for DENV reverse genetics as described above. The *in vitro* transcribed DENV RNA library was electroporated in either Vero or C6/36 cells, the viral supernatants were passaged 3 times every 4 to 5 days in the corresponding cells for enrichment.

### High-throughput sequencing and analysis

Viral RNA was isolated using a QIAamp Viral RNA Mini Kit (Qiagen). Amplicons containing the library regions were prepared for sequencing through two rounds of PCR, using the Illumina TruSeq system and Q5 Hot Start DNA polymerase (NEB). Primers for the first round of PCR were specific to the DENV2 prM sequence with overhangs for Illumina adapters. This PCR product was purified and used as a template for a second round of PCR using the standard Illumina P5 and P7 primers with barcodes and sequencing adaptors. PCR products were purified and analyzed on a Qubit 4 fluorometer (Invitrogen) and Bioanalyzer (Agilent Technologies) for quality control. Amplicon libraries were diluted to 4 nM and pooled for sequencing, which was carried out on a MiSeq system with 300bp paired-end reads. Plasmid and P0 libraries were sequenced at a depth of ∼1 million reads per sample; further passages were sequenced with depth between 300,000 – 1 million reads to sample. A custom perl script^44^ was used to analyze the sequences, and custom R scripts were used to plot the data.

### Furin cleavage prediction

Furin cleavage site efficiency was predicted using the Pi-Tou software^25^, providing amino acids from position P14-P6’ of the DENV furin cleavage sites.

### Statistical analysis

Statistical analysis was carried out using Graphpad Prism version 9.0. Growth kinetics and maturation of DENV variants were compared to their corresponding wildtype using 2-way ANOVA multiple comparisons. Neutralization titers of DENV variants were compared to their corresponding wildtype using Student’s t-test. Significance symbols are defined as follow: * p<0.05, ** p< 0.01, *** p<0.001, **** p<0.0001. Data are graphed as mean +/-standard deviation.

## Acknowledgments

We thank members of the Baric and de Silva laboratories for helpful discussions. This project received support from NIH grants AI107731 and AI125198 to A.M.D and R.S.B. P01 AI106695 to A.M.D.. L.V.T. is the recipient of the Pfizer NCBiotech Distinguished Postdoctoral Fellowship in Gene Therapy.

## Author Contributions

L.V.T and R.S.B designed the study. R.M.M perform high-throughput sequencing preparation and analysis. L.V.T., R.M.M., S.D., L.E.A., L.J.W. performed experiments. L.V.T., A.M.D., R.S.B. provide oversight of the project. L.V.T. wrote the manuscript. R.M.M and R.S.B. reviewed and revised the final version.

## Conflict of Interest

L.V.T. R.M.M. and R.S.B. are inventors on a patent application filed on the subject matter of this manuscript.

---

**Supplementary Figure 1:**
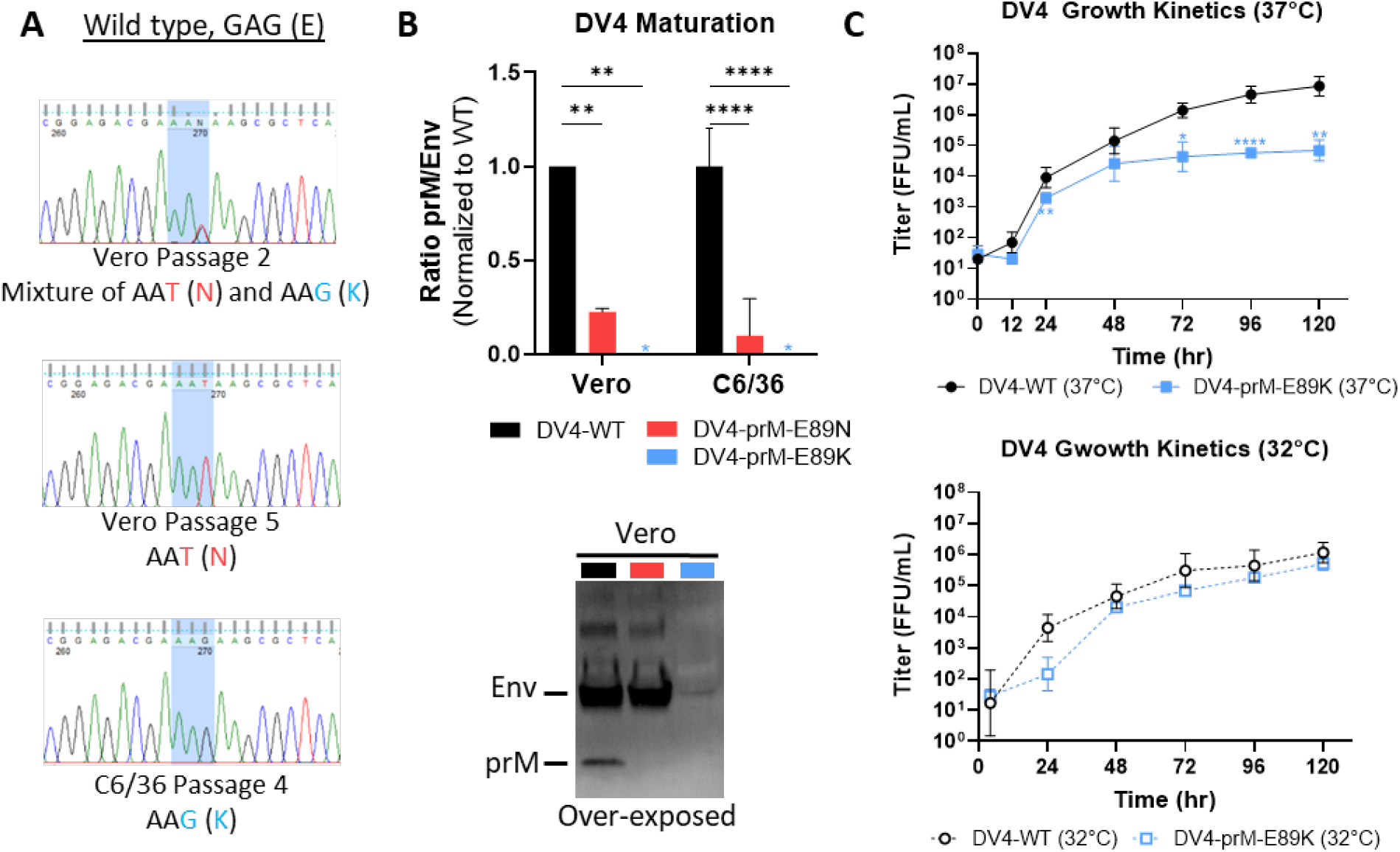
(A) DNA Chromatograms of DV4-E89K in C6/36 cells (bottom) as well as Vero cells from early (P2, top) and late (P5, middle) passage. (B) Representative western blot image (bottom) of DV4-WT, DV4-prM-E89K and DV4-prM-E89N viral supernatant blotted with anti-Env and anti-prM antibodies, and quantification of viral maturation (prM/Env) normalized to DV4-WT. (C) Growth kinetics of DV4-WT and DV4-prM-E89K in Vero and C6/36 cells at 32°C (bottom) and 37°C (top).

**Supplementary Figure 2:**
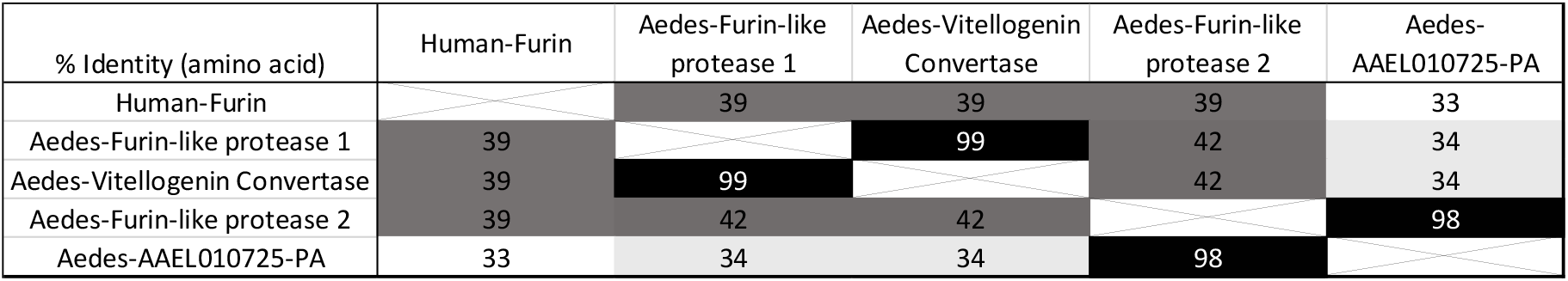
Amino acid sequence identity matrix of furin and furin-like proteases between human and *Aedes aegypti*.

**Supplemental Table 1:**
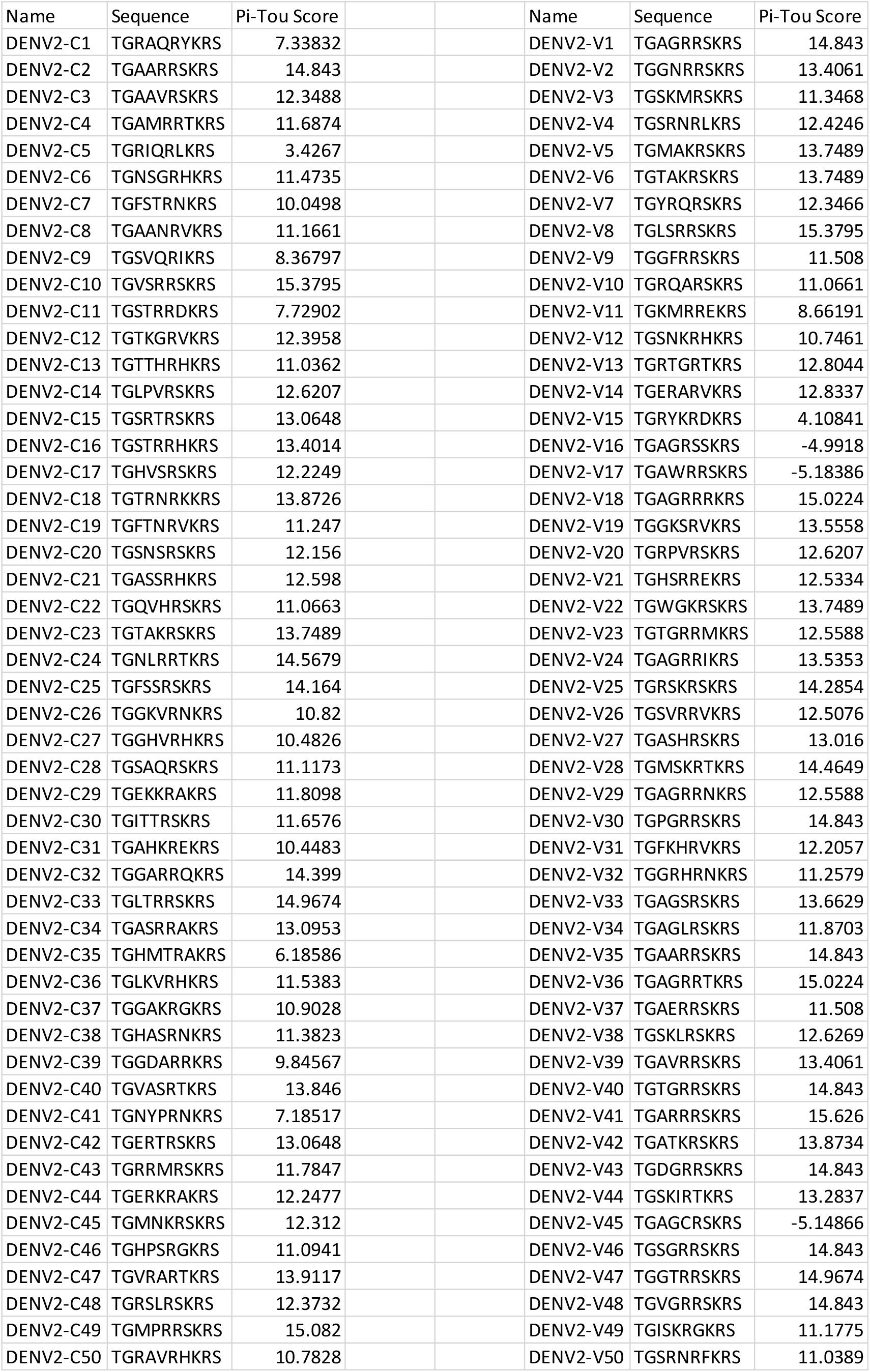
Top 50 enriched sequences and PiTou scores of DV2 directed-evolution.

**Supplemental Table 2:**
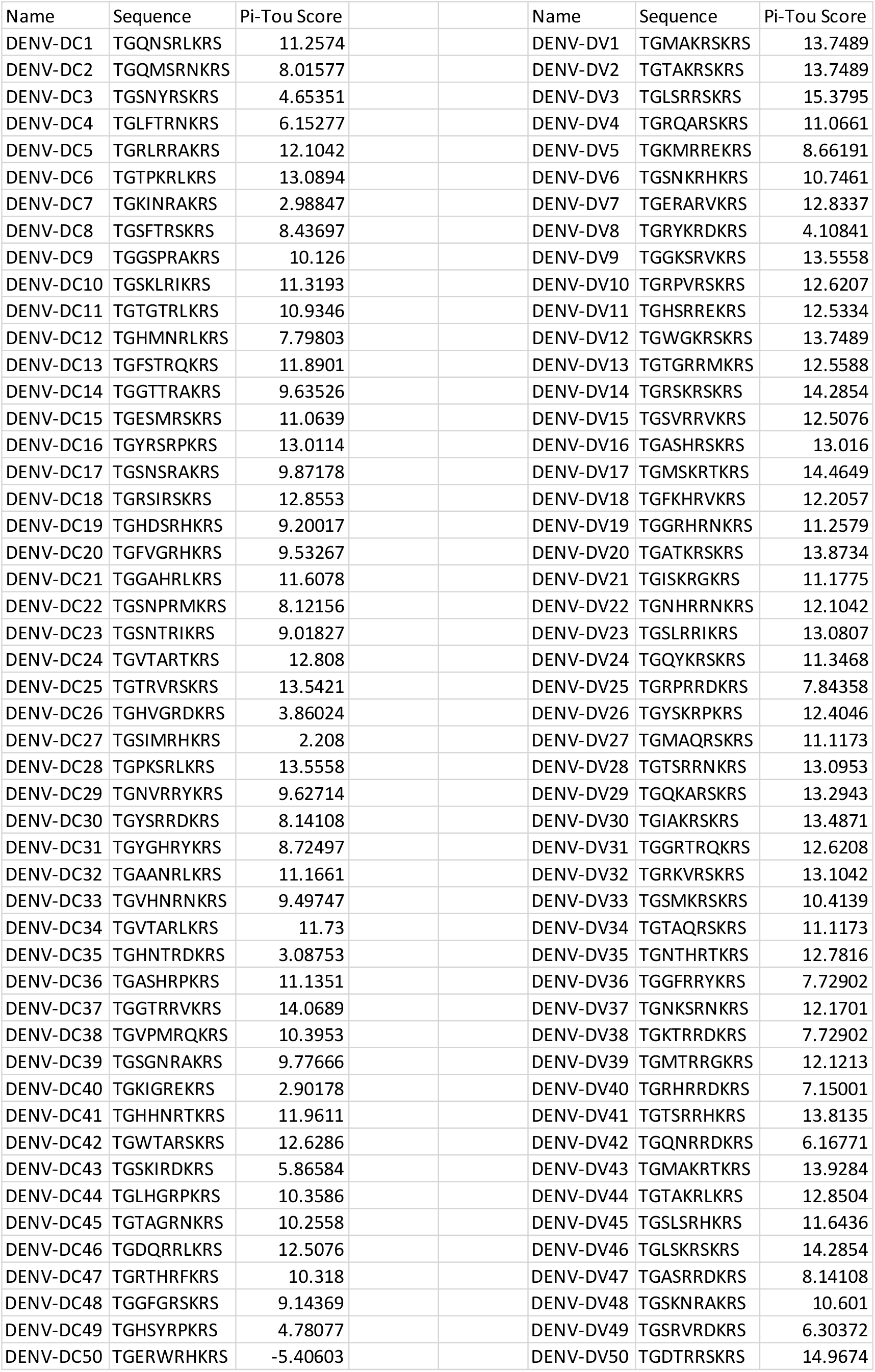
Top 50 deselected sequences and PiTou scores of DV2 directed-evolution.

## Notes

### Competing Interest Statement

The authors have declared no competing interest.

